# Contributions of error correction and the spindle assembly checkpoint to mitotic timing and fidelity

**DOI:** 10.64898/2026.03.10.710927

**Authors:** Gloria Ha, Luyi Qiu, Ariel Amir, Daniel J. Needleman

## Abstract

Chromosome segregation is a tightly-regulated process that normally occurs with high fidelity. Errors in chromosome segregation are associated with aging, cancer, and infertility. Initially erroneously attached chromosomes are corrected over the course of mitosis, with the spindle assembly checkpoint preventing entry into anaphase until this error correction is complete. Despite extensive work on the molecular basis of error correction and the spindle assembly checkpoint, it is still unclear how disruption of these processes contribute to chromosome segregation errors. Here, we develop and experimentally test a coarse-grained model of error correction in the presence of a faulty spindle assembly checkpoint. We use the resulting model to disentangle the impact of various small molecule and genetic perturbations on both error correction and the spindle assembly checkpoint, and to compare chromosomally stable hTERT-RPE-1 cells and chromosomally unstable U2-OS cells. We find that the probability of error-free chromosome segregation is determined by the ratio of the checkpoint failure rate to the error correction rate, and validate a simple heuristic for understanding the source of chromosome segregation errors: perturbations which cause errors by disrupting the spindle assembly checkpoint decrease anaphase times, while those that disrupt error correction increase anaphase times. Taken together, this work provides a quantitative framework for understanding how error correction and the spindle assembly checkpoint contribute to mitotic timing and fidelity.

## Introduction

Chromosomes are segregated by the spindle during cell division. When the spindle functions properly, the chromosomes are equally divided into the two daughter cells. Chromosome segregation errors (i.e. non-equal division of chromosomes) are strongly associated with cancer [1–3] and are the primary cause of infertility and pregnancy loss [4–6]. The rate of chromosome segregation errors varies widely between systems and circumstances: from below 0.001% in Saccharomyces cerevisiae [7], to ≈ 0.1% in tissue culture cells [8], to above 10% in human oocytes [9]. Despite extensive work, it remains unclear what precise mechanisms underlie these differences or how they produce the associated quantitative variations in chromosome segregation error rates.

Chromosome segregation errors are suppressed by a combination of two processes: error correction and the spindle assembly checkpoint. Initially erroneous attachments between chromosomes and the spindle are corrected over the course of spindle assembly [10–13]. The spindle assembly checkpoint detects erroneous attachments and prevents entry into anaphase until it no longer senses their presence [14–16]. While there have been many advances in uncovering the molecular machinery responsible for error correction and the spindle assembly checkpoint, it is less well understood how they work as systems, or how their dysfunctions impact chromosome segregation errors.

We previously developed a procedure to characterize the dynamics of error correction in mitosis [17]. We used measurements of the number of chromosomes in the two daughter cells after the mother cell was prematurely forced into anaphase, as well as measurements of the spontaneous anaphase times, both of which could be quantitatively explained by a simple model of error correction. In that model, initially erroneously attached chromosomes are autonomously corrected at a constant rate, with the cell transitioning to anaphase after the last erroneously attached chromosome is corrected. That prior work only addressed circumstances in which the spindle assembly checkpoint was well functioning. In the present study, we sought to expand this work to include conditions with faulty spindle assembly checkpoints. We find that a model in which a faulty checkpoint fails at a constant rate is inconsistent with the measured anaphase times, while a model in which the rate of checkpoint failure depends on the number of erroneous attachments can quantitatively explain the distribution of anaphase times and the distribution of the number of chromosomes in the subsequent daughter cells. We use the resulting combined model of error correction in the presence of an imperfect spindle assembly checkpoint to disentangle the quantitative effects of a variety of molecular perturbations, and to compare chromosomally stable hTERT RPE-1 cells and chromosomally unstable U2-OS cells. Analytical analysis of the model provides insight into trends observed across the different conditions. A key finding is that the probability of error-free chromosome segregation is determined by the ratio of the checkpoint failure rate to the error correction rate. Our work also supports the use of a simple heuristic to determine which process is disrupted by a perturbation that increases segregation errors: if the perturbation increases anaphase times, then it primarily impacted error correction; if the perturbation decreased anaphase times, then it primarily impacted the spindle assembly checkpoint. Taken together, this work leads to a quantitative framework for understanding how error correction and the spindle assembly checkpoint contribute to mitotic timing and fidelity.

## Results

### Perturbing the spindle assembly checkpoint decreases anaphase onset times and increases chromosome segregation errors

We previously established a coarse-grained model of error correction in cells with a well-functioning spindle assembly checkpoint [17]. In this model, an erroneously attached chromosome is defined to be one in which both chromatids would partition into the same daughter cell upon entry to anaphase. There are on average *C*_*E*,init_ initially erroneouslyattached chromosomes at the beginning of spindle assembly. It is assumed that error correction is a chromosome-autonomous process, with erroneously attached chromosomes corrected at rate *k*_*b*_ and correctly attached chromosomes becoming erroneous at rate *k*_*e*_ (Fig. 1A). If the spindle assembly checkpoint is functioning perfectly, then the cell will only enter anaphase after the last chromosome is corrected, resulting in an equal number of chromatids in the two daughter cells: i.e. the difference in the number of kinetochores between the daughter cells, |∆*N* |, will be zero. The distribution of anaphase times in this model can be solved analytically, and agrees with measurements of chromosomally stable hTERT-RPE-1 human tissue culture cells, which, as expected, show |∆*N* | = 0 for nearly all pairs of daughter cells (Fig. 1B). Forcing hTERT-RPE-1 into anaphase early with an Mps1 inhibitor results in highly elevated values of |∆*N* |, which decreases with time in mitosis in a manner consistent with the predictions of the coarse-grained model (Fig. 1C). The agreement between observations and the model ‘s prediction provide support for the model ‘s underlying assumptions, such as chromosome autonomous error correction, and simultaneously fitting the spontaneous anaphase times and forced anaphase data, enables measurements of the models parameters, including the rate of error correction: *k*_*b*_ = 0.55 ± 0.02 min^−1^ in hTERT-RPE-1 cells.

**FIG. 1:**
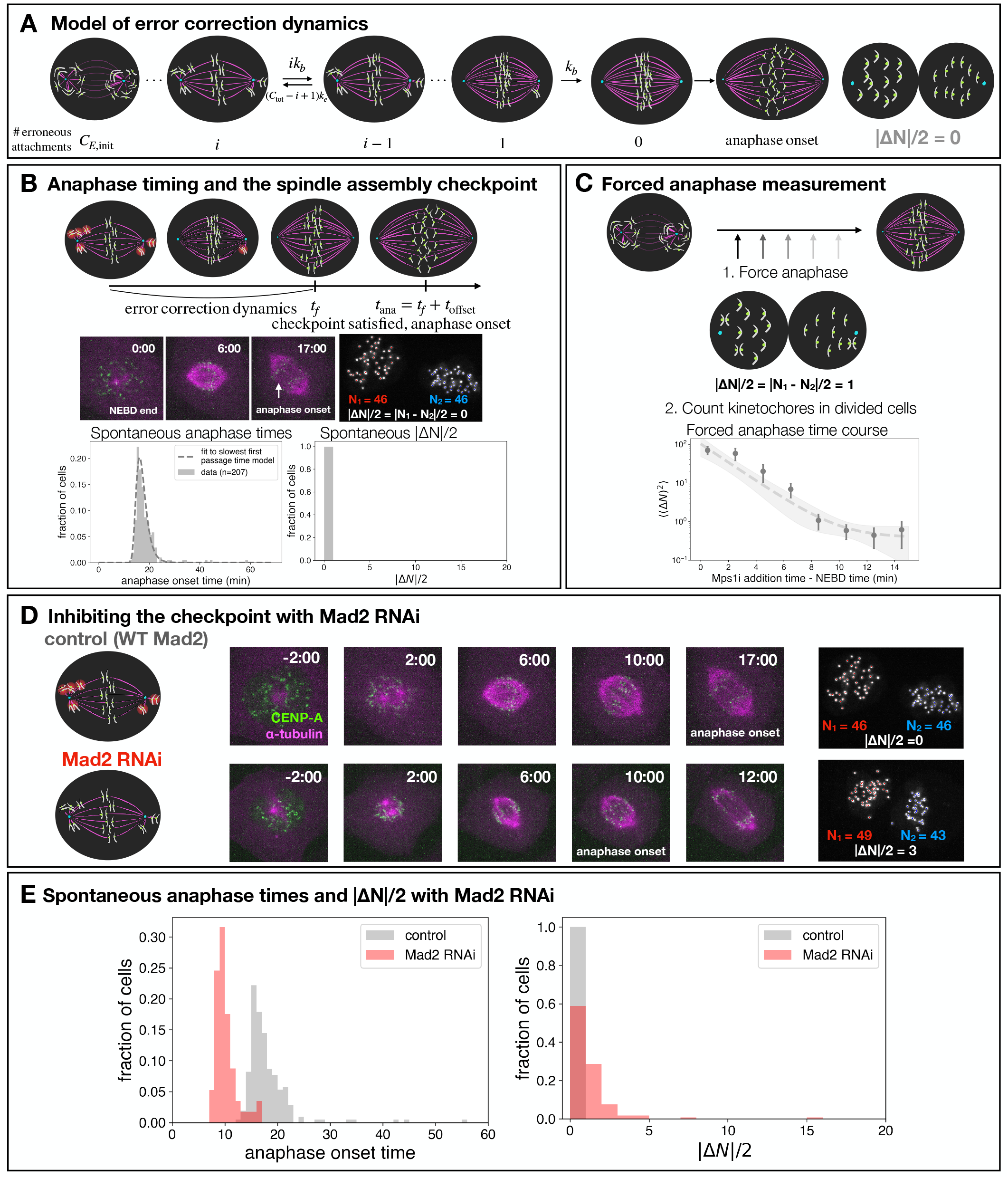
Perturbing the spindle assembly checkpoint decreases anaphase onset times and increases chromosome segregation errors. (A) Schematic of the coarse-grained model of error correction with a perfect spindle assembly checkpoint (SAC). Anaphase occurs after the last erroneous chromosome is corrected, resulting in a kinetochore count difference, |∆*N* |*/*2, of 0 between daughter cells. (B) The onset of anaphase occurs at a time *t*_offset_ after the time, *t*_*f*_, at which the last erroneously attached chromosome is corrected (upper). Images of cells during mitosis and high resolution images of the kinetochores in the daughter cells (middle), allows measurements of the distributions of both anaphase times and a spontaneous |∆*N* |*/*2 (lower, grey), and comparison with model predictions (lower, dotted lines). (C) Forcing anaphase prematurely using an Mps1 inhibitor at different times relative to nuclear envelope breakdown (NEBD) and subsequent measurements of the kinetochore count differences in the daughter cells (lower, circles) enables comparison with model predictions (lower, dashed line) and characterization of error correction dynamics.(D) Comparison of control (wild type MAD2) and Mad2 RNAi example images from time-lapse microscopy (center) and kinetochore counts after division (right). (E) Resulting spontaneous anaphase times and |∆*N* |*/*2 distributions without (grey) and with (red) Mad2 knocked down via RNAi.

If the spindle assembly checkpoint always behaved as ideally conceptualized in this model, then spontaneous chromosome segregation errors would never occur. However, spontaneous errors do occur under numerous conditions [18, 19]. For example, using RNA interference to knockdown Mad2, which localizes to erroneously-attached kinetochores and generates a “wait anaphase” signal [20–22](Fig. 1D), results in reduced anaphase times [23] and increased |∆*N* | values between daughter cells, indicating increased spontaneous errors [17](Fig. 1E). We thus sought to generalize this coarse-grained model to better understand conditions with imperfect spindle assembly checkpoints and spontaneous chromosome segregation errors.

### State-dependent model best recapitulates Mad2 RNAi data

Since Mad2 plays a central role in the spindle assembly checkpoint but is not believed to contribute to error correction, explaining the impact of Mad2 knockdown provides a particularly simple test case for models of error correction in the presences of a faulty spindle assembly checkpoint. To construct such a model, we used the previously-defined coarse-grained dynamics of chromosomes autonomous error correction (Fig. 1A): a spindle in a state with *i* erroneous attachments proceeds to state *i* − 1 at rate *k*_*b*_*i* and to state *i* + 1 at rate *k*_*e*_(*C*_tot_ − *i*), where *C*_tot_ is the total number of chromosomes. In the case of a perfect spindle assembly checkpoint, the cell progresses to anaphase after the last erroneous chromosome is corrected, i.e. after it enters the *i* = 0 state. In the case of a faulty spindle assembly checkpoint, the transition to anaphase does not only occur from the *i* = 0 state, but also from other states which still have erroneous attachments, i.e. from *i* ≠ 0 states.

We first considered a model in which spindles enter anaphase with a constant “checkpoint failure rate”, *k*_*f*_, independently of the number of their erroneous attachments (Fig. 2A). Using this constant rate model of a faulty spindle assembly checkpoint, and assuming an equal probability of erroneous attachments being associated with either pole, we numerically calculate the predicted distribution of anaphase onset times and the predicted distributions of |∆*N* |*/*2 after anaphase, both of which depend on *k*_*f*_ (Appendix). For *k*_*f*_ = 0, this model is equivalent to the previously described perfectly function spindle assembly checkpoint model (Fig. 1A) [17]. For *k*_*f*_ ≠ 0, two populations of cells appear in the anaphase time distributions (Fig. 2B): one population arises from cells entering anaphase with rate *k*_*f*_, i.e. from states *i* ≠ 0, and thus exhibit an exponentially decaying distribution of anaphase times. The other population results from cells entering anaphase after all of their chromosomes correctly attach, i.e. from state *i* = 0, and exhibit a peaked distribution of anaphase times. For higher *k*_*f*_, the population that enters anaphase prematurely dominates the population that enters anaphase after correcting all attachments. The constant rate model also predicts an increasingly broad distribution of spontaneous |∆*N* |*/*2 as *k*_*f*_ increases (Fig. 2B).

**FIG. 2:**
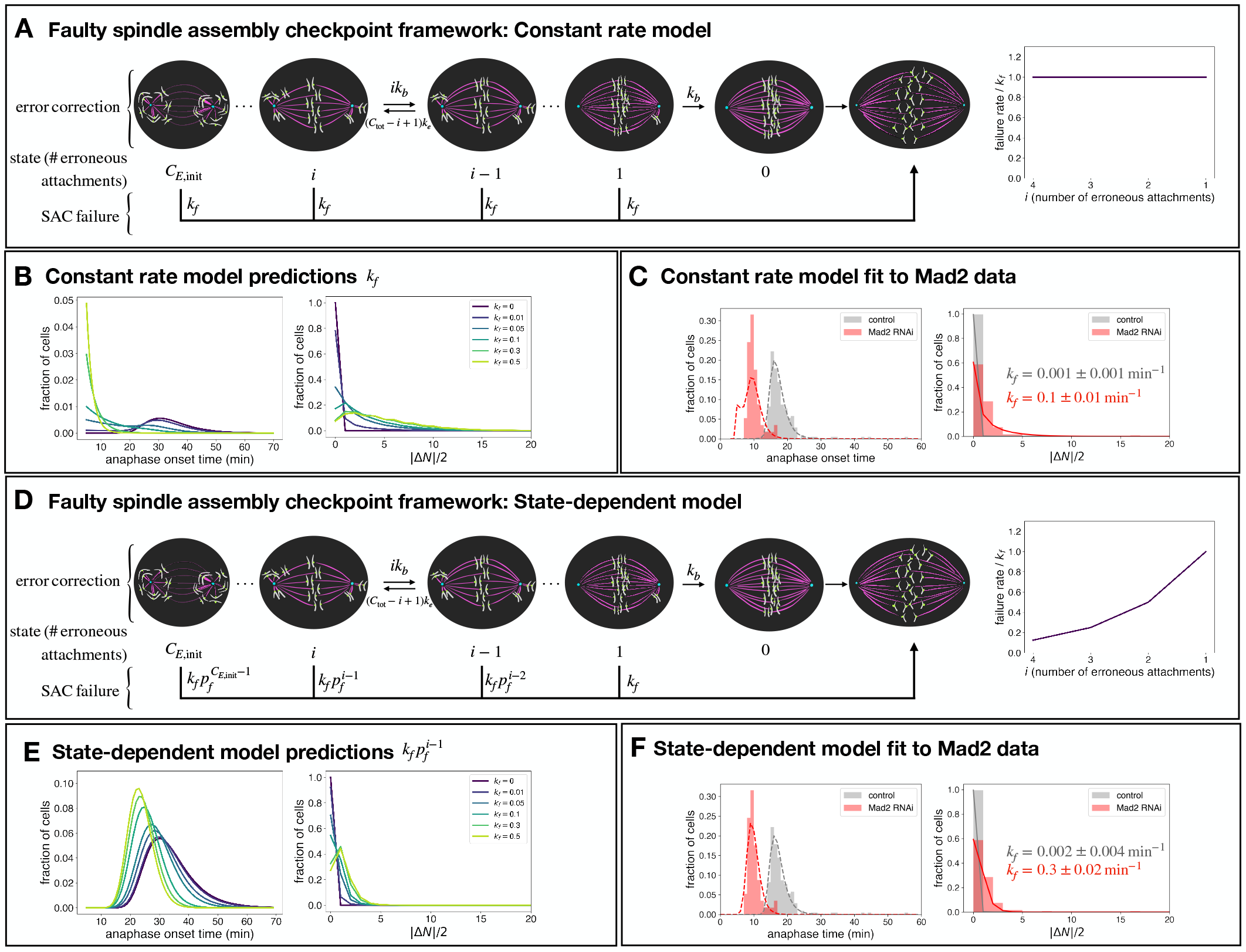
Models of a faulty spindle assembly checkpoint. (A) Schematic of the constant rate model of spindle assembly checkpoint failure, in which the rate of a faulty transition to anaphase is a constant, *k*_*f*_, and thus independent of the number of erroneous attachments. (B) Predicted anaphase time distributions and kinetochore count distributions under the constant rate model for different values of *k*_*f*_ . (C) Constant rate model fit (lines) results for anaphase time and kinetochore count distributions, compared to experimental observations (shaded bars), without and with Mad2 RNAi. (D) Schematic of the state-dependent model of spindle assembly checkpoint failure, in which the rate of a faulty transition to anaphase increases with decreasing numbers of erroneous attachments. (E) Predicted anaphase time distributions and kinetochore count distributions under the constant rate model for different values of *k*_*f*_ . (F) State-dependent model fit results for anaphase time and kinetochore count distributions without (gray) and with (red) Mad2 RNAi.

We next sought to compare the predictions of the constant rate model with the results from the Mad2 RNAi experiments. Since Mad2 does not play a role in error correction, we assumed that the rates of error correction *k*_*b*_, error formation *k*_*e*_, an the initial number of errors *C*_*E*,init_, had the same value in Mad2 RNAi as those previously determined for controls (0.55±0.02 min^−1^, 0.001±0.0002 min^−1^, and 26±3 respectively) [17]. We then simultaneously fit the anaphase time distribution and spontaneous |∆*N* |*/*2 distributions measured for Mad2 RNAi to the predictions of the constant faulty checkpoint model (Fig. 2C), and found clear discrepancies: Due to the non-zero *k*_*f*_ required by the wide |∆*N* |*/*2 distribution in the Mad2 RNAi data, the constant rate model predicts that far more cells should enter anaphase early (less than 7 minutes) than are actually observed to.

The inadequacy of the constant rate model, which is predicated on the assumption that the rate of checkpoint failure is independent of the number of the erroneous attachments, inspired us to construct an alternative model in which the checkpoint failure rate does depend on spindle state. We considered a state-dependent model in which the rate of checkpoint failure decreases rapidly with the number of erroneous attachments, *i*, as 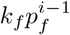 (Fig. 2D). One rationale for this functional form is that if the probability for a single erroneous attachment to escape checkpoint is *p*_*f*_, then in the presence of *i* erroneous attachments the checkpoint failure rate should be proportional to 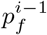. In this model, the probability of faulty anaphase becomes vanishingly small as the number of erroneous attachments increases. While the constant rate model predicts a bimodal distribution of anaphase times, which is not observed experimentally (Fig. 2B), the state-dependent model predicts a unimodal anaphase time distribution, that decreases in mean and variance with increasing *k*_*f*_, as well as a distributions of spontaneous |∆*N* |*/*2 which becomes progressively wider as *k*_*f*_ increases (Fig. 2E).

We then used the state-dependent model to fit both the Mad2 RNAi and control experimental data (Fig. 2F). The fits sensitively depended on the value of *k*_*f*_, but, while *p*_*f*_ = 0.5 gave the best fit to the Mad2 RNAi data, values of *p*_*f*_ between 0.1 and 0.6 yielded similar results (Fig. S1), so, for simplicity, we fixed *p*_*f*_ = 0.5 and fit for *k*_*f*_ . We found that the best-fit *k*_*f*_ increased dramatically from *k*_*f*_ = 0.002 ± 0.004 min^−1^ for the control data to *k*_*f*_ = 0.3 ± 0.02 min^−1^ for the Mad2 RNAi data. The fits to both the anaphase time and the spontaneous |∆*N* |*/*2 results were excellent, indicating that the state-dependent model can quantitatively describe mitotic behaviors in the presence of a faulty spindle assembly checkpoint (Fig. 2F).

### Faulty SAC model can recapitulate small molecule perturbation results

We next sought to leverage the state-dependent faulty SAC model to investigate the effects of small molecule perturbations on both error correction and SAC function. We used the previously described washout procedure of the kinesin-5 inhibitor monastrol [17] as the baseline “control” condition for these experiments. Monastrol exposed spindles exhibit increased numbers of initially erroneous attachments, which can be advantageous for studying error correction [17, 24, 25] and cells arrested in metaphase through monastrol incubation are easily identified and enter mitosis synchronously upon drug washout, which aids in data collection. We synchronized hTERT RPE-1 cells using 2 hours of incubation in monastrol, leading to the formation of monopolar spindles. We then washed out the monastrol during imaging, leading to spindle bipolarization and anaphase (Fig. 3A). Monastrol washout in RPE-1 cells slightly increases the baseline error rate (from 0.5% non-zero |∆*N* |*/*2 in the RPE-1 control case to 4% non-zero |∆*N* |*/*2 in the monastrol washout case), but still provides a low baseline with which to compare perturbations – when we simultaneously fit the |∆*N* |*/*2 distribution, anaphase time distribution, and forced anaphase kinetochore count data, the best-fit checkpoint failure rate was *k*_*f*_ = 0.005 ± 0.001 min^−1^, which is in the same order of magnitude as the RPE-1 control condition (*k*_*f*_ = 0.002 ± 0.004 min^−1^, Fig. 3B).

**FIG. 3:**
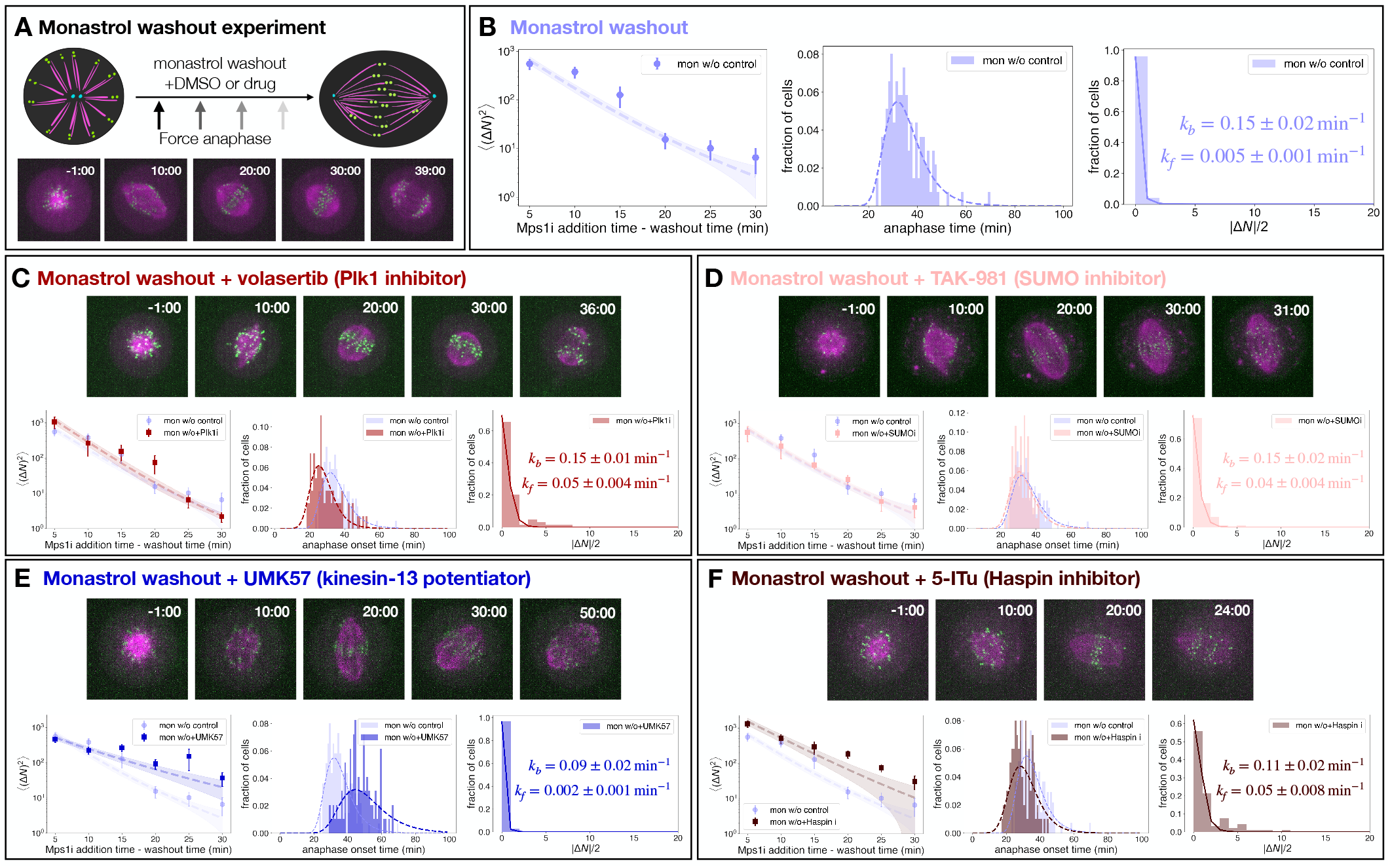
Small molecule perturbations of error correction and the spindle assembly checkpoint. (A) Forced anaphase time course, anaphase onset time distribution, and |∆*N* |*/*2 distribution for monastrol washout 0.5% v/v DMSO control data. Data was simultaneously fit to the full state-dependent faulty SAC model. (B) Forced anaphase time course, anaphase onset time distribution, and |∆*N* |*/*2 distribution for monastrol washout with Plk1 inhibitor (30nM volasertib) data. Anaphase times are faster and spontaneous |∆*N* |*/*2 values are higher, but the error correction rate is indistinguishable from control. (C) Forced anaphase time course, anaphase onset time distribution, and |∆*N* |*/*2 distribution for monastrol washout with SUMO-activating enzyme inhibitor (10uM TAK-981) data. Spontaneous |∆*N* |*/*2 values are higher, but the error correction rate is indistinguishable from control. (D) Forced anaphase time course, anaphase onset time distribution, and |∆*N* |*/*2 distribution for monastrol washout with kinesin-13 potentiator (1uM UMK57) data. Anaphase times are slower and the error correction rate is slower, but the spontaneous |∆*N* |*/*2 values are indistinguishable from control. (E) Forced anaphase time course, anaphase onset time distribution, and |∆*N* |*/*2 distribution for monastrol washout with Haspin inhibitor (1 µM 5-ITu) data. Error correction rate is slower, but the spontaneous |∆*N* |*/*2 values are higher, leading to slightly faster anaphase times than control.

We first targeted Plk1, a kinase that localizes to prometaphase kinetochores, stabilizes correct kinetochoremicrotubule attachments, and regulates the spindle assembly checkpoint through recruitment of Mad2 to unattached kinetochores [26, 27]. We inhibited Plk1 using volasertib, a small molecule inhibitor of Plk1[28]. Volasertib has previously been studied for clinical applications – when added at high concentrations to interphase cells, volasertib induces monopolar spindle arrest and reduces cell proliferation [28, 29].

When we added 30nM of volasertib during the washout from monopolar arrest, cells formed bipolarized spindles and divided (Fig. 3C). Volasertib-treated cells entered anaphase 29 ± 1 minutes after release from the monopolar initial state (*n* = 81 cells), which was significantly faster than in the DMSO control (35 ± 1 minutes). Additionally, the baseline |∆*N* |*/*2 was elevated from the control – 34% of volasertib-treated cells had a non-zero kinetochore count difference (*n* = 64 cells), compared to 4% in the DMSO control. We performed a simultaneous fit on the |∆*N* |*/*2 distribution, anaphase time distribution, and forced anaphase timecourse data. We found that the best fit error correction rate for the volasertib data was *k*_*b*_ = 0.15 ± 0.01 min^−1^ and that the checkpoint failure rate was *k*_*f*_ = 0.05 ± 0.004 min^−1^ (Fig. 3C). Thus Plk1 inhibition through volasertib treatment results in an unchanged error correction rate from the monastrol washout control, and a significantly increased checkpoint failure rate.

We next targeted the small ubiquitin-like modifier (SUMO) enzymatic cascade [30]. SUMOylation is a posttranslational modification that has been found to regulate the localization of many spindle proteins including Aurora B kinase and kinetochore components [31]. We used TAK-981, a small molecule inhibitor of the SUMO-activating enzyme that has been shown to induce severe mitotic dysfunction and polyploidy over long term incubation [32, 33].

Cells treated with (10 µM) of TAK-981 upon monopolar washout divided in 31 ± 0.5 minutes on average (*n* = 94 cells), which was faster than the monastrol washout control (35 ± 1 minutes). The baseline |∆*N* |*/*2 was also elevated from 4% in the DMSO control to 26% in the TAK-981 condition (*n* = 85 cells), indicating a faulty spindle assembly checkpoint. We simultaneously fit the spontaneous |∆*N* |*/*2 distribution, anaphase time distribution, and forced anaphase data using the state-dependent faulty checkpoint model, and found that the best fit error correction rate *k*_*b*_ was 0.15 ± 0.02 min^−1^, which was within error identical to the monastrol washout control (0.15 ± 0.02 min^−1^). The initial error state was also indistinguishable to the DMSO control – ⟨(∆*N* )^2^⟩(*t* = 5 min) = 553 ± 253 for TAK-981 treated cells, vs. 558 ± 142 for the monastrol washout control, The checkpoint failure rate, however, increased from *k*_*f*_ = 0.005 ± 0.001 min^−1^ in the monastrol washout control to *k*_*f*_ = 0.04 ± 0.004 min^−1^ in the TAK-981 treated cells. In summary, inhibition of SUMOylation through TAK-981 resulted in an indistinguishable initial error state and error correction rate compared to the monastrol washout control, but had a higher SAC failure rate *k*_*f*_, resulting in faster anaphase times and a higher rate of non-zero spontaneous |∆*N* |*/*2.

We then used UMK57, a small molecule potentiator of kinesin-13 motors, to test the effect of activating MCAK during monastrol washout [34]. Since kinesin-13 motors regulate microtubule detachment from the kinetochore, activating kinesin-13 through UMK57 treatment leads to microtubule destabilization. We previously used UMK57 to activate MCAK in RPE-1 cells, and saw that high concentrations (1 µM) of UMK57 led to both slower anaphase times and slower error correction compared to the control [17]. Adding 1 µM UMK57 to RPE-1 cells during washout from monastrol incubation led to a significant increase in mean anaphase time from 35 ± 1 minutes in the monastrol washout control to 46 ± 9 minutes with UMK57. The kinetochore count difference distribution was similar to the control, with only 2.8% of cells having a non-zero kinetochore count difference with UMK57, compared to 4% in the monastrol washout control (Fig. 3E). When we simultaneously fit the data, we found that the error correction rate *k*_*b*_ was significantly decreased from *k*_*b*_ = 0.15 ± 0.02 min^−1^ in the monastrol washout control to *k*_*b*_ = 0.09 ± 0.02 min^−1^ with UMK57. The checkpoint failure rate of *k*_*f*_ = 0.002 ± 0.001 min^−1^ with UMK57 was similar to the monastrol washout control. Similar to the experiments in unsynchronized cells, activating MCAK through UMK57 decreased the error correction rate and slowed down anaphase times without affecting the spindle assembly checkpoint.

We then tested the effects of 5-ITu, a small molecule inhibitor of Haspin kinase, which is an important regulator of error correction [35, 36]. Haspin kinase localizes to erroneously attached kinetochores and recruits Aurora B kinase, which in turn destabilizes microtubule attachments and promotes error correction [37, 38]. Adding 1 µM 5-ITu to RPE-1 cells during washout from monastrol incubation shifted the the anaphase time distribution, with the mean anaphase time changing from 35 ± 1 minutes in the monastrol washout control to 28 ± 6 minutes with 5-ITu. The initial state of ⟨(∆*N* )^2^⟩ was significantly increased, from 560 ± 140 in the monastrol washout control to ⟨(∆*N* )^2^⟩(*t* = 5 min) = 1300 ± 570 with 5-ITu, indicating higher statistical asymmetry in the initial configuration of attachments under Haspin inhibition. Haspin inhibition also led to a lower error correction rate of *k*_*b*_ = 0.11 ± 0.02 min^−1^. Finally, the checkpoint failure rate was increased to *k*_*f*_ = 0.05 ± 0.008 min^−1^ under Haspin inhibition. Haspin inhibition led to a higher initial state of errors, slower error correction rate, faster anaphase times, and higher checkpoint failure rate.

In summary, these perturbations of Plk1, SUMOylation, MCAK, and Haspin kinase, demonstrate that the statedependent faulty checkpoint model is consistent with experimental results obtained under a variety of conditions, and can be used to quantitatively distinguish perturbations that affect the initial error state, error correction rate, and spindle assembly checkpoint fidelity.

### Characterizing ploidy and errors in U2-OS cancer cells

The perturbation and control experiments described above were performed using the euploid, chromosomally stable cell line, hTERT-RPE1, and were consistent with the state-dependent faulty spindle assembly checkpoint model. We next sought to determine if the state-dependent faulty spindle assembly checkpoint model could also capture the behaviors of a cell line with chromosomal instability, and thus be used to extract the initial error state, the rate of error correction, and the degree of SAC fidelity in such cell lines. We chose to use U2-OS, an near-tetraploid osteosarcoma-derived human cell line commonly used as a model cancer cell line. We generated a stable U2-OS cell line expressing CENP-A::sfGFP, mCherry::alpha tubulin, and emiRFP670::hCentrin2 using a lentiviral system. We stained the cells with SPY650-DNA to identify the timing of chromosome condensation and nuclear envelope breakdown (Fig. 4A).

**FIG. 4:**
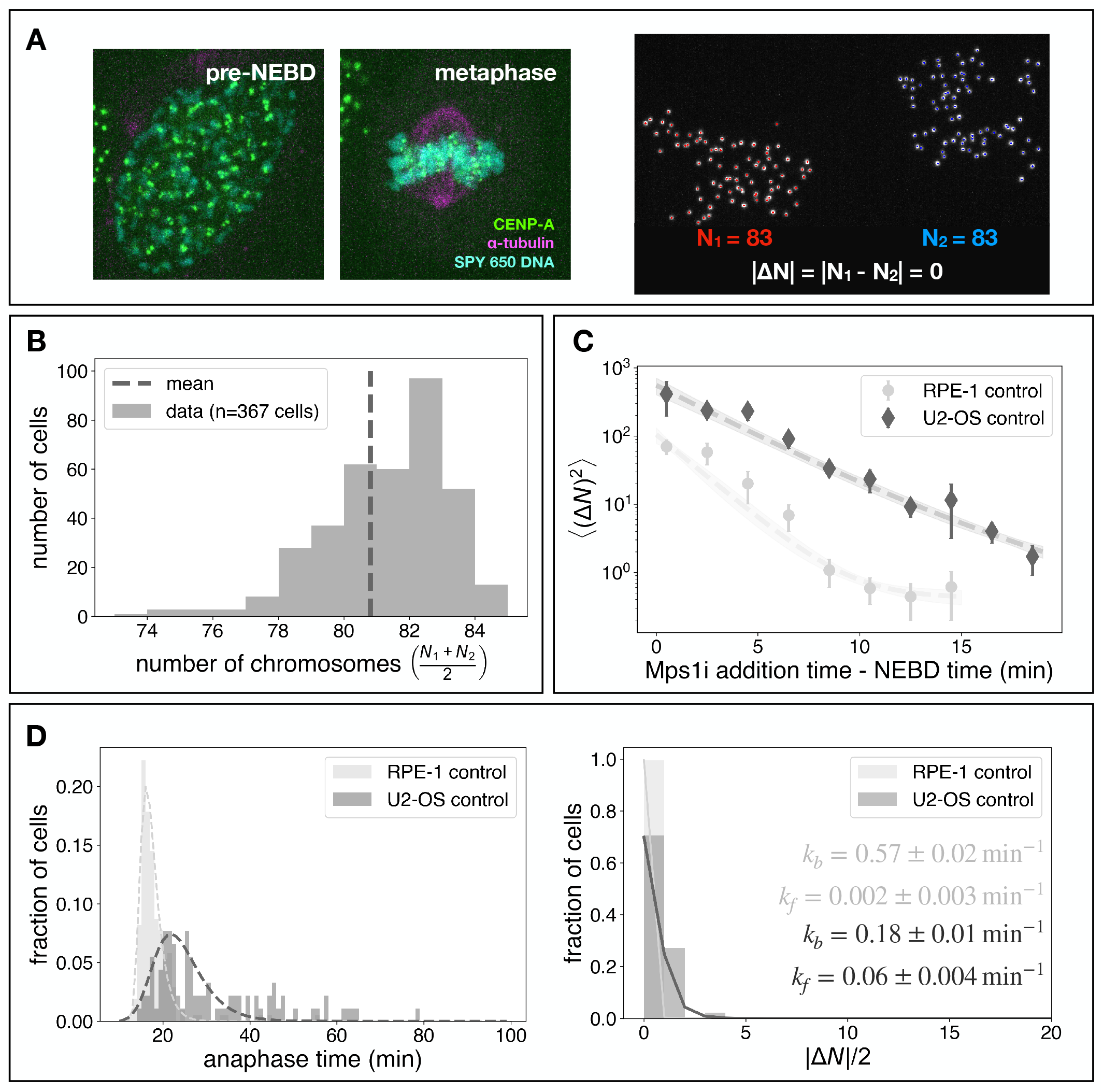
Characterizing ploidy, error correction dynamics, and SAC function in U2-OS cells. (A) Example of a U2-OS cell before nuclear envelope breakdown, in metaphase, and after division, when kinetochore counts are measured. (B) Chromosome count distribution in U2-OS cells as measured by the sum of kinetochores in the divided daughter cells. (C) Error correction comparison between control RPE-1 and U2-OS cells. (D) Anaphase onset time distribution and |∆*N* |*/*2 distribution for control RPE-1 and U2-OS cells. All model fits shown are from simultaneously fitting all 3 datasets to the state-dependent faulty checkpoint model.

We first sought to characterize the ploidy of our U2-OS cell line. By counting the total number of kinetochores in the divided daughter cells, we measured ploidies ranging from 73 to 84 chromosomes, with an average of 80.8 ± 0.1 chromosomes per cell (*n* = 367 cells, Fig. 4B), similar to plodies of 65 to 80 chromosomes previously reported in U2-OS cell [39, 40]. We next investigated error correction dynamics, the distribution of anaphase times, and the spontaneous error rate in this cell line. We used the forced anaphase assay to characterize error correction dynamics and found that ⟨(∆*N* )^2^⟩ were higher in U2-OS cells than in RPE-1 cells and appeared to decay more slowly with time, indicating a higher initial error state and slower error correction dynamics in U2-OS cells (Fig. 4C). We measured the spontaneous anaphase onset times in these U2-OS cells and found that they were both more variable and slower than RPE-1 cells (Fig. 4D, left), with a mean anaphase time of 31 ± 1 minutes on average (*n* = 91 cells, time measured from nuclear envelope breakdown to kinetochore separation), compared to 17.6 ± 0.3 minutes in RPE-1 cells. The spontaneous|∆*N* |*/*2 values were higher in U2-OS than in RPE-1 cells (Fig. 4D, right), with 29% of U2-OS cells exibiting a non-zero |∆*N* |*/*2 (*n* = 92 cells) compared to 0.5% in RPE-1 cells, indicating a faulty spindle assembly checkpoint in U2-OS cells. We used the state-dependent model of spindle assembly checkpoint failure to simultaneously fit the forced anaphase data, the distribution of spontaneous anaphase times, and the distribution of spontaneous |∆*N* |*/*2 values (Fig. 4C,D). The resulting rate of error correction in U2-OS cells was *k*_*b*_ = 0.18 ± 0.01 min^−1^, substantially slower than in RPE-1 cells (*k*_*b*_ = 0.57 ± 0.02min^−1^), while the best-fit spindle assembly checkpoint failure rate for U2-OS cells was *k*_*f*_ = 0.06 ±0.004 min^−1^, significantly higher than the failure rate for RPE-1 cells (*k*_*f*_ = 0.002 ±0.003 min^−1^). Thus, the state-dependent faulty checkpoint model can describe the behaviors of both euploid, chromosomally stable RPE-1 cells and aneuploid, chromosomally unstable U2-OS cells, and can be used to quantify their differences in error correction and spindle assembly checkpoint fidelity.

### Perturbations of initial state and error correction differentially affect error correction in U2-OS cells

We next sought to characterize error correction and the spindle assembly checkpoint in U2-OS cells subject to perturbations. We first targeted kinesin-13 motor activity using the kinesin-13 activator, UMK57. UMK57 in low doses has been shown to decrease the incidence of lagging chromosomes in cancer cells, elderly cells, and embryos [34, 41, 42]. In particular, short-term (*<* 1 hour) incubation with 100 nM UMK57 has been shown to decrease the percentage of U2-OS cells with lagging chromosome from 32% to 15% [34].

We added 100 nM UMK57 media to U2-OS cells immediately before imaging, resulting in imaged cells being exposed to UMK57 for between 5 to 30 minutes before entering mitosis (Fig. 5A). Forced anaphase experiments revealed similar ⟨(∆*N* )^2^⟩ time courses for UMK57-treated and control U2-OS cells (Fig. 5B). UMK57-treated U2-OS cells also exhibited similar distributions of spontaneous anaphase times and similar distributions of |∆*N* |*/*2 values as control U2-OS cells (Fig. 5B). Simultaneously fitting all three sets of data and resulted in indistinguishable error correction rates (*k*_*b*_ = 0.17 ± 0.02 min^−1^ with UMK57, *k*_*b*_ = 0.18 ± 0.01 min^−1^ without UMK57) and spindle assembly failure rates (*k*_*f*_ = 0.08 ± 0.002 min^−1^ with UMK57, *k*_*f*_ = 0.06 ± 0.004 min^−1^ without UMK57) for UMK57-treated and control U2-OS cells. Taken together, these results indicate that a dose and duration of UMK57 exposure that was previously reported to decrease lagging chromosomes in U2-OS cells did not significantly impact error correction rate or spindle assembly failure rate.

**FIG. 5:**
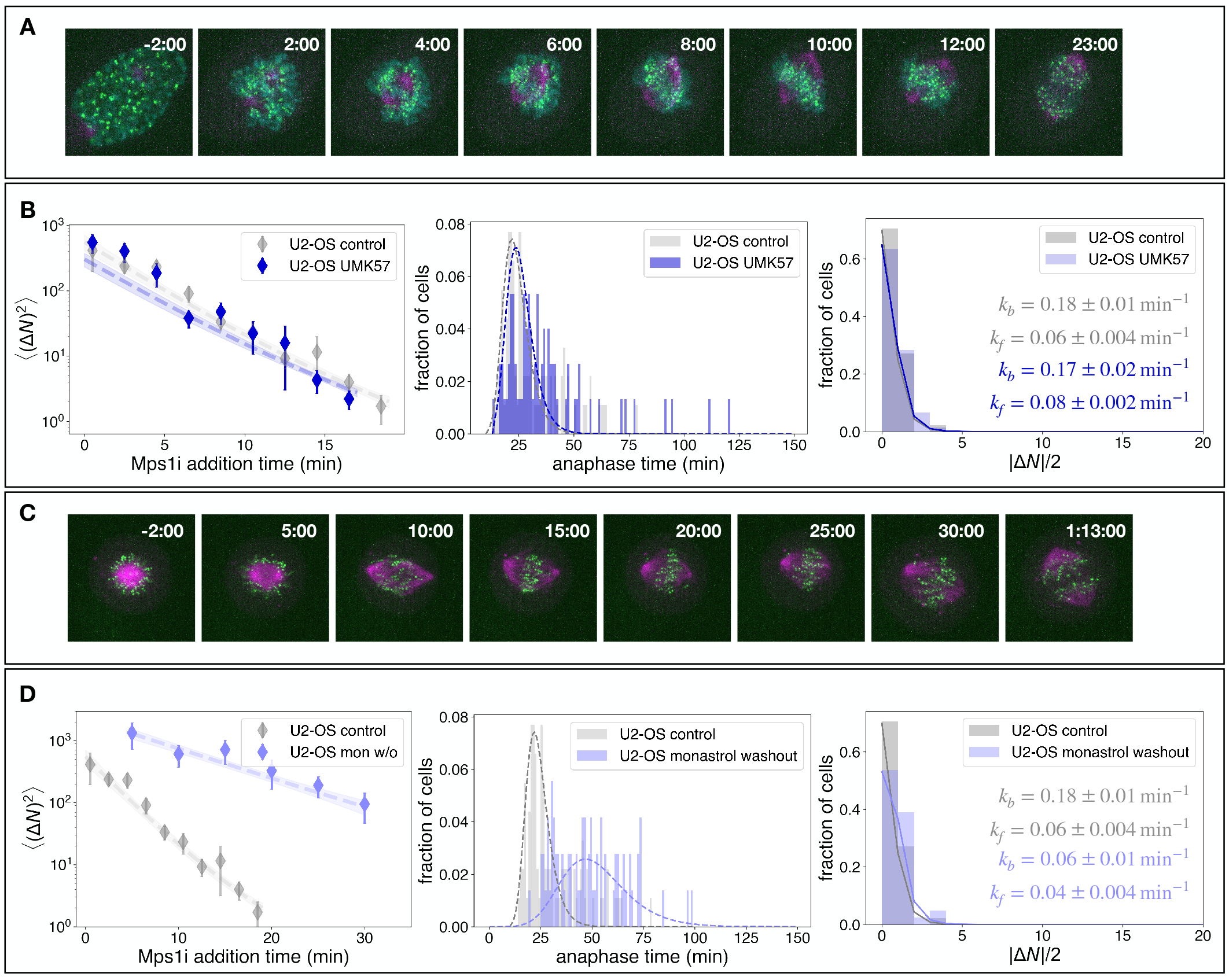
Perturbations of initial state and error correction differentially affect error correction in U2-OS cells. (A) Example of unsynchronized U2-OS cell undergoing mitosis with 100nM UMK57 (kinesin-13 potentiator). (B) Forced anaphase time course, anaphase onset time, and spontaneous |∆*N* |*/*2 data for U2-OS cells with 100nM UMK57. (C) Example of U2-OS cell released from monastrol, leading to spindle bipolarization and eventually division. (D) Forced anaphase time course, anaphase onset time, and spontaneous |∆*N* |*/*2 data for U2-OS cells undergoing monastrol washout.

We next tested the effect of the initial configuration of attachments on error correction in U2-OS cells. We once again used monastrol, a kinesin-5 motor inhibitor that arrests spindles in a monopolar state [43]. We incubated U2-OS cells in 100 µM monastrol for 2.5 hours and then imaged the cells as we washed out monastrol, allowing for spindle bipolarization and division (Fig. 5C). Forced anaphase experiments revealed that ⟨(∆*N* )^2^⟩ in the monastrol washout U2-OS cells were significantly higher than controls for all time lags (Fig. 5D), similar to the impact of monastrol on RPE-1 cells [17] Fig. 3B). In the monastrol washout, U2-OS cells divided significantly slower than controls, going from a mean anaphase time of 31 ± 1 minutes in the control (*n* = 91 cells) to 50 ± 2 minutes in the monastrol washout cells (*n* = 72 cells), similar to the shift observed in RPE-1 cells with and without monastrol washout [17] Fig. 3B). Additionally, the frequency of non-zero spontaneous |∆*N* |*/*2 values increased from 29% in the control (*n* = 92 cells) to 46% in the monastrol washout U2-OS cells (*n* = 41 cells). Simultaneous fitting all three datasets gave an error correction rate of *k*_*b*_ = 0.06 ±0.01 min^−1^ for monastrol washout U2-OS cells, significantly less than control U2-OS cells (*k*_*b*_ = 0.18 ± 0.01 min^−1^), while the best fit spindle assembly failure rate *k*_*f*_ was similar between the two conditions, with *k*_*f*_ = 0.06 ± 0.004 min^−1^ for the control and *k*_*f*_ = 0.04 ± 0.004 min^−1^ for monastrol washout. Thus, in both RPE-1 cells and U2-OS cells, starting from an initially monopolar state leads to a substantial decrease in the rate of error correction without impacting spindle assembly checkpoint fidelity.

### Spindle assembly checkpoint failure effect can be approximated by the last step

Our results described above indicate that the state-dependent faulty checkpoint model is consistent with data from two different cell lines subject to multiple molecular perturbations, and numerical fitting with this model enables quantitative measurements of the error correction rate and checkpoint failure rate. We next sought to investigate trends across the different conditions. To do so, we began by further analyzing the state-dependent faulty checkpoint model to obtain insight into the expected trends.

In the presence of a perfect spindle assembly checkpoint (*k*_*f*_ = 0), anaphase commences a time *t*_offset_ after the last erroneously attached chromosome is corrected (Fig. 1A,B). Thus, calculating the distribution of anaphase times with a perfectly function spindle assembly checkpoint is a slowest first passage time problem. In the (relevant) limit that the error correction rate, *k*_*b*_, is much larger than the rate of generation of new errors, *k*_*e*_, the anaphase times follow a Gumbel distribution with mean anaphase time [17]:

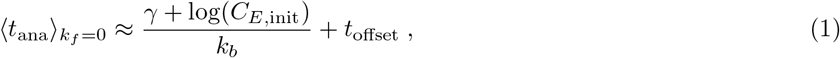

where *γ* ≈ 0.5772 is Euler ‘s constant.

To extend such analytical results to cases in which the checkpoint is faulty (*k*_*f*_ ≠ 0), we note in the Mad2 RNAi data (Fig. 1E), where one of the most central components of the checkpoint is being directly perturbed, most cells have a spontaneous kinetochore count difference |∆*N* |*/*2 = 0 (59%), followed by |∆*N* |*/*2 = 1 (29%). Since, even in this extreme case, 88% of cells have spontaneous |∆*N* |*/*2 ≤ 1, we hypothesized that the major contribution to faulty anaphase comes from cells dividing with one chromosome segregation error. We formalize this hypothesis by making a “last step approximation”, in which a cell dividing with *i* = 1 error is the dominant form of faulty anaphase. In this case, the mean anaphase time can be expressed as the sum of the mean first passage time to reach state *i* = 1 and the expected time to go from state *i* = 1 to division. From state *i* = 1, the cell can proceed to division via two different paths: dividing in the presence of one error at rate *k*_*f*_ or correcting the last error at rate *k*_*b*_ before dividing (Fig. 6A). When the checkpoint is perfect (*k*_*f*_ = 0) the expected time to go from state *i* = 1 to division is 1*/k*_*b*_. When there is a non-zero rate *k*_*f*_ for the system to divide at state *i* = 1, the total rate for the system to go from state 1 to division is *k*_*b*_ + *k*_*f*_ and the expected time to division becomes 1*/*(*k*_*b*_ + *k*_*f*_ ). Thus the difference in the mean anaphase time induced by faulty spindle assembly checkpoint in the last step is simply 1*/*(*k*_*b*_ + *k*_*f*_ ) − 1*/k*_*b*_, yielding the following equation for the approximate mean anaphase time in the presence of a faulty checkpoint:

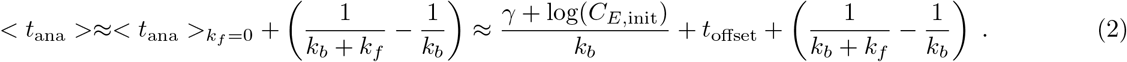

**FIG. 6:**
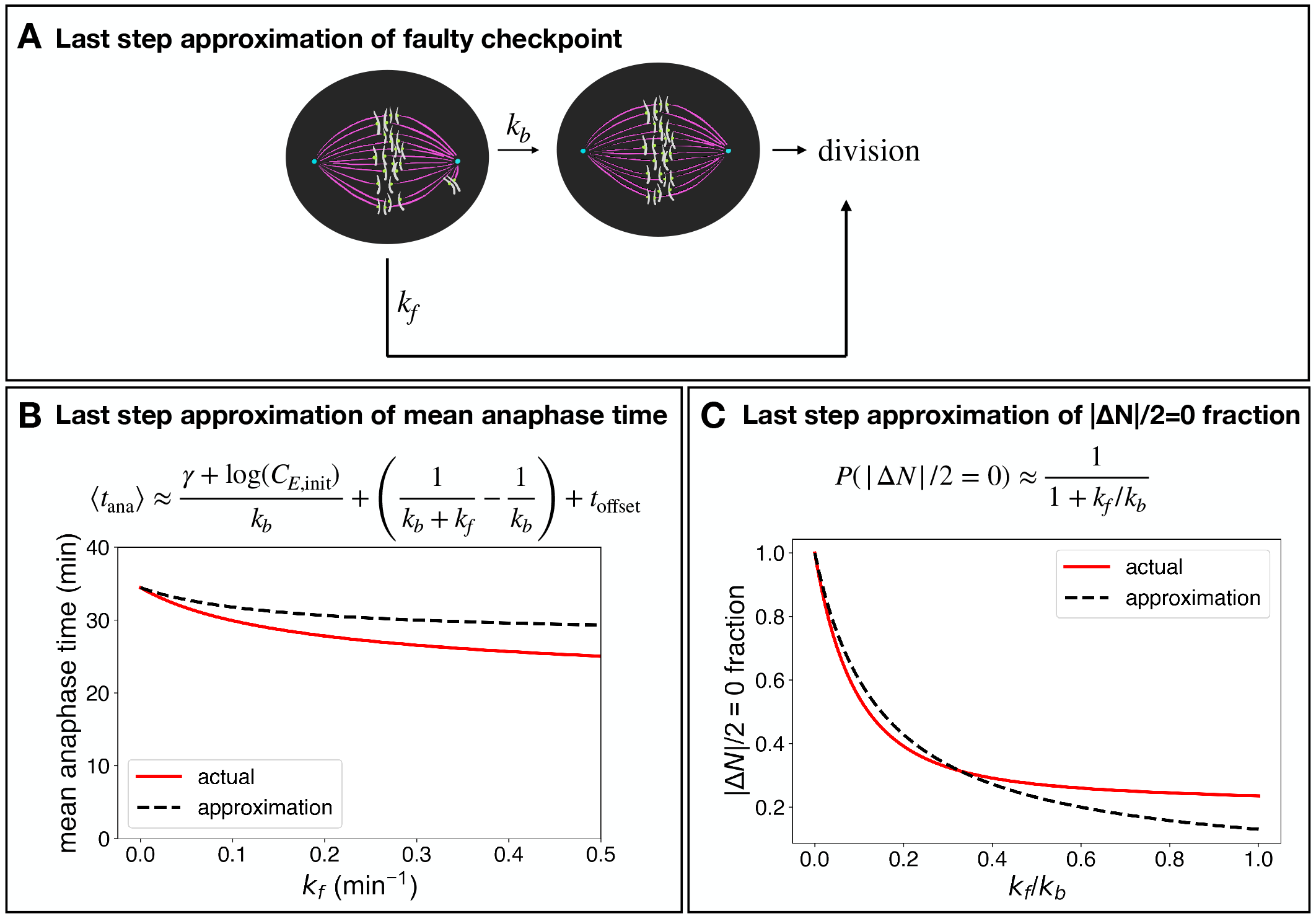
Spindle assembly checkpoint failure effect can be approximated by the last step. (A) Schematic of last step approximation. (B) Approximate mean of anaphase times considering only leakage with 1 error. Fixed *k*_*b*_ = 0.15 min^−1^ and *p*_*f*_ = 0.5. (C) Approximate |∆*N* |*/*2 = 0 fraction only considering the contribution of dividing with one error. Fixed *k*_*b*_ = 0.15 min^−1^ and *p*_*f*_ = 0.5.

Thus, as intuitively expected, the mean anaphase time decreases as either error correction rate *k*_*b*_ or checkpoint failure rate *k*_*f*_ increases. When we compare the last step approximation to the numerically calculated mean anaphase time of the full state-dependent faulty assembly checkpoint model, we see that the approximation is within 10% of the actual value for the relevant range of *k*_*f*_ (Fig. 6B).

We next investigated predictions for the spontaneous kinetochore count difference using the same last step approximation. If we assume that cells can divide with either 0 errors (after correcting at rate *k*_*b*_) or 1 error (at rate *k*_*f*_ ), the fraction of cells that divide with 0 errors and have |∆*N* |*/*2 = 0 is given by the ratio of these rates:

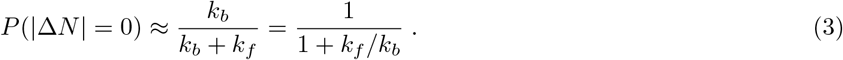

We compared the predictions of Eq. 3 with the numerical results of the full state-dependent faulty assembly checkpoint model (Fig. 6C), and found that the approximation reproduced the relationship between the kinetochore count differences and the rates of checkpoint failure and error correction. However, for higher values of *k*_*f*_ */k*_*b*_, predictions from the last step approximation were systematically higher than those from the full model. When *k*_*f*_ is higher, the last step approximation over estimates the expected *P* (|∆*N* | = 0) as more cells can divide with more than one error. A key insight from this analysis is that the fraction of cells with the correct number of chromosomes after division, i.e. the fraction of cells with |∆*N* | = 0, depends on the *ratio* of checkpoint failure rate to the error correction rate, not just on the checkpoint failure rate itself. This highlights that a perturbation that enhances chromosome segregation error can do so by increasing the rate of checkpoint failure, decreasing that rate of error correction, or both.

Eq. 2 relates the model ‘s parameters to the mean anaphase time, and Eq. 3 relates the model ‘s parameters to the fraction of cells with the correct number of chromosomes after division. These two equations can be combined to give further insight into the impact of perturbations. Consider a molecular perturbation that only changed the checkpoint failure rate, causing it to go from 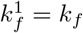, before the perturbation, to 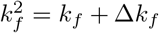, after the perturbation. Use of Eq. 2 and Eq. 3 show that this would result in a change in both anaphase times, 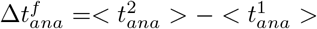, and a change in fraction of cells with the correct number of chromosomes after division, ∆*P*^*f*^ (|∆*N* | = 0) = *P* ^2^(|∆*N* | = 0) − *P* ^1^(|∆*N* | = 0), with

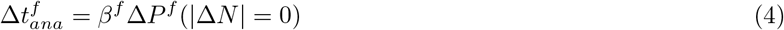

Where *β*^*f*^ = 1*/k*_*b*_, the constant of proportionality relating 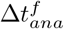 and ∆*P*^*f*^ (|∆*N* | = 0), is positive. Thus, Eq.4 indicates that if the perturbation increases the checkpoint failure rate, it will result in a decrease in both the fraction of cells with the correct number of chromosomes after division and the mean anaphase time.

A similar analysis demonstrates that a perturbation that only changed the error correction rate, causing it to go from 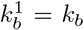, before the perturbation, to 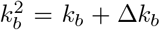, would result in a change in both anaphase times, 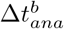, and the fraction of cells with the correct number of chromosomes after division, ∆*P*^*b*^(|∆*N* | = 0), with

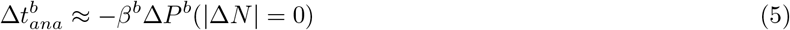

Where 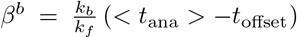, the constant of proportionality relating 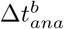 and ∆*P*^*b*^(|∆*N* | = 0), is positive. Thus, Eq.5 indicates that if the perturbation decreases the error correction rate, it will result in an decrease in the fraction of cells with the correct number of chromosomes after division and an increase in the mean anaphase time.

### Perturbation results can be described by the last step approximation

We sought to use the results from the last step approximation to gain insights into the trends across the different conditions we investigated. The analysis presented in Fig 2, Fig 3, Fig 4, and Fig 5, as well as additional experiments with UMK57 [17] and inhibition of Aurora B (see appendix), resulted in measurement of error correction rates and checkpoint failure rates in both RPE-1 and U2-OS cells under a variety of conditions, with *k*_*f*_ varying over one hundred fold (from 0.002/min to 0.3/min) and *k*_*b*_ varying nearly ten fold (from 0.06/min to 0.57/min) Fig. 7A. The results from all these conditions collapse onto one master curve when the spontaneous |∆*N* |*/*2 = 0 fraction is plotted versus *k*_*f*_ */k*_*b*_, quantitatively consistent with the prediction from the last-step approximation (Fig. 7B). This result indicates the utility of the last-step approximation and emphasizes that chromosome segregation errors result from the interplay between error correction and checkpoint failure. Furthermore, the agreement between measurements and the predictions of the last step approximation indicate that an estimate of *k*_*f*_ */k*_*b*_ could be obtained by simply measuring the fraction of cells with the correct number of chromosome after division, i.e. the spontaneous |∆*N* |*/*2 = 0 fraction, without needing to perform the forced anaphase experiments or measurements of the distribution of anaphase times.

**FIG. 7:**
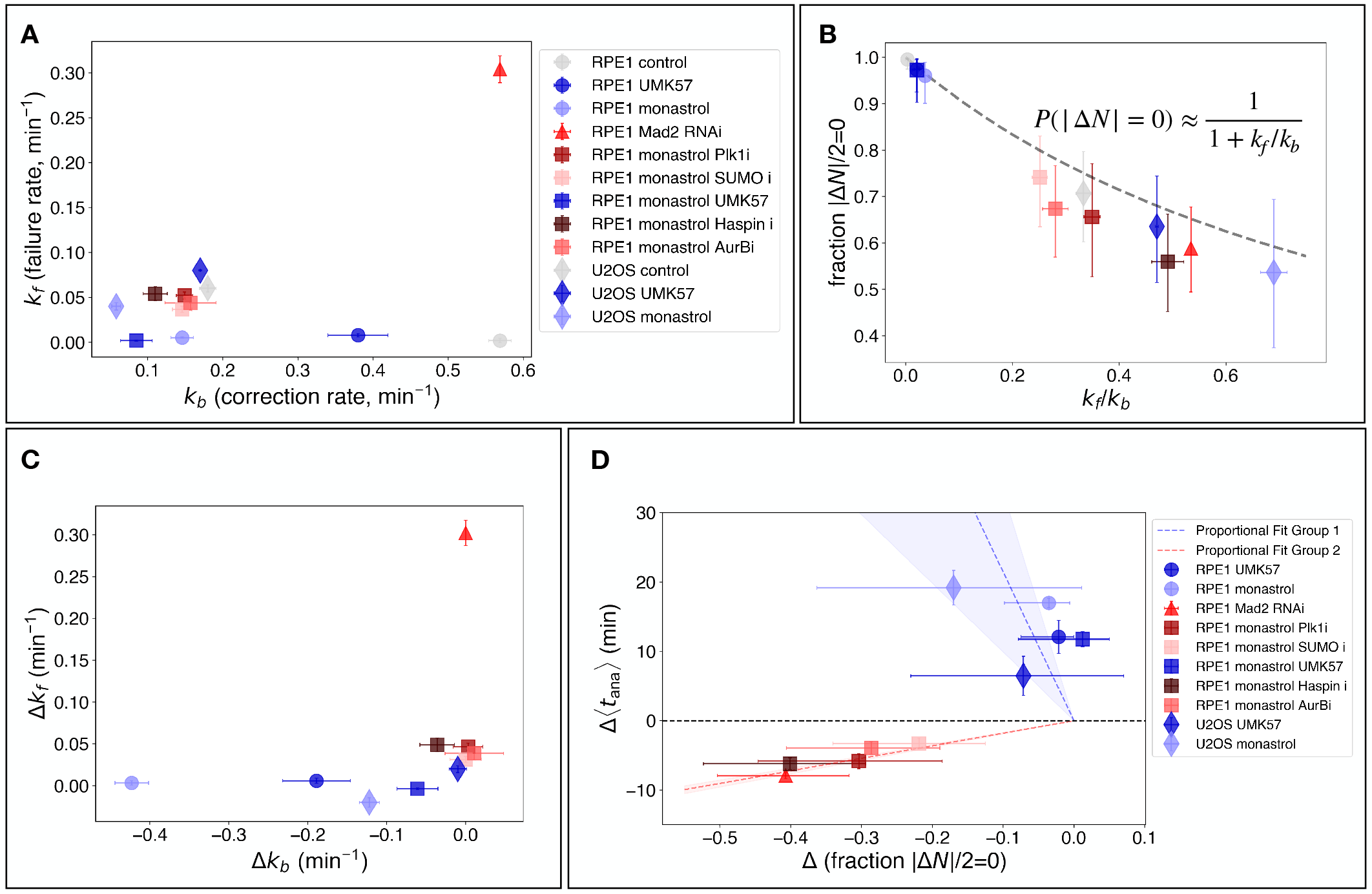
Summary of mitotic perturbations in RPE-1 and U2-OS cells. (A) Comparison of spindle assembly checkpoint failure rate *k*_*f*_ vs. error correction rate *k*_*b*_ across conditions. (B) The fraction of cells with |∆*N* |*/*2 = 0 across conditions are consistent with predictions from the last step approximation, which yields a simple dependence on the ratio of rates *k*_*f*_ */k*_*b*_ (dashed line). (C) Perturbations can be divided into two classes: those which primarily affect the error correction rate *k*_*b*_ (blue) and those which primarily affect the checkpoint failure rate *k*_*f*_ (red). Change in parameter values calculated relative to relevant control conditions (see main text). (D) The change in mean anaphase time and the change in the fraction of cells with |∆*N* |*/*2 = 0 are proportional to each other, with a positive slope for perturbations which primarily impact the checkpoint failure rate (red, the best-fit slope is 18 ± 1 min) and a negative slope for perturbations which primarily impact the error correction rate (blue, the best-fit slope is −220 ± −120 min).

We next investigated how these different perturbations altered the rates of error correction and checkpoint failure. To do so, we calculate the differences in *k*_*f*_ and *k*_*b*_ between the different conditions and a corresponding control, where for single perturbations the control used was the baseline RPE1 or U2-OS values (as appropriate) and where for dual drug conditions involving both monastrol and another drug, the values with only monastrol present served as the control. The perturbations could be split into two groups: (i) perturbations which impacted the rate of checkpoint failure more than the rate of error correction (red, Fig. 7C), inhibition of Mad2, Plk1, SUMO, Haspin, and Aurora B; (ii) perturbations which impacted the rate of error correction more than the rate of checkpoint failure (blue, Fig. 7C), UMK57 and monastrol applied either singly or together to either RPE1 cells or U2-OS cells. The group of perturbations that primarily impacted the rates of checkpoint failure exhibit decreases in both the fraction of cells with the correct number of chromosomes after division and mean anaphase times (red, Fig. 7D). The group of perturbations that primarily impacted the rates of error correction exhibit decreases in the fraction of cells with the correct number of chromosomes after division and increases in mean anaphase times (blue, Fig. 7D). These observed trends are consistent with those anticipated from Eq.4 and Eq.5. For both groups, the changes in mean anaphase times are approximately proportional to the changes in the fraction of cells with the correct number of chromosomes after division (dotted lines, Fig. 7D), though the fitted values of the slopes are only roughly similar to those calculated using all the measured model parameters (6.2 ± 2.7 min vs fitted 18 ± 1 min for the group in which checkpoint failure rates are primarily impacted, and −800 ± 900 min vs fitted −220 ± 120 min for the group in which error correction rates are primarily impacted). Eq.4 and Eq.5 were derived under the assumption that perturbations impact only *k*_*f*_ or *k*_*b*_, while multiple parameters actually do change across conditions, so it is not surprising that they do not qualitatively capture the observations. Nonetheless, Eq.4 and Eq.5 do semi-qualitatively explain the trends in the data. This suggests that measurements of mean anaphase time and the fraction of cells with the correct number of chromosomes after division can be used to infer whether a perturbation predominantly affects error correction or checkpoint fidelity, without needing to perform the forced anaphase experiment or fit data to the full state-dependent faulty checkpoint model.

## Discussion

In this study, we investigated the behaviors of spindles with a faulty spindle assembly checkpoint. We used forced anaphase experiments, and measurements of the distribution of spontaneous anaphase times and the distribution of kinetochore counts in the daughter cells, to characterize conditions with chromosomal instability, including both genetic and small molecule perturbations and cancer cells. All of these experimental results can be quantitatively explained using a coarse-grained model in which the rate of checkpoint failure depends on the number of erroneous kinetochore attachments. A key finding, consistent with both experimental observations and analytical calculations, is that the probability of error-free chromosome segregation is determined by the ratio of the checkpoint failure rate to the error correction rate. Taken together, this work provides a framework for understanding how the interplay between perturbations to error correction and the spindle assembly checkpoint impact the fidelity of chromosome segregation.

Previous mechanistic models of the spindle assembly checkpoint have focused on modeling the molecular basis of its behaviors [44–47]. The coarse-grained model developed here provides a means of studying the consequences of checkpoint failure, including the impact on experimentally-measured anaphase time distributions and on spontaneous segregation errors, as measured by the distribution of |∆*N* |*/*2 kinetochore count differences between daughter cells. we incorporated spindle assembly checkpoint failure as an “escape rate” from any state of *i* erroneously attached chromosomes. A model in which the escape rate is a constant, irrespective of the state of attachments, is inconsistent with the data. An alternative, state-dependent model in which the escape rate strongly decreases with the number of erroneous attachments, can quantitatively explain our experimental results for all conditions investigated. In cells with a properly functioning checkpoint, a single incorrectly attached chromosome is sufficient to activate the spindle assembly checkpoint and block anaphase onset [21]. In cells with a faulty checkpoint, a decrease in escape rate with increasing number of erroneous attachments might result from a weakened, but still present, “wait anaphase” signal being emitted from erroneously attached chromosomes, consistent with previous work suggesting that the checkpoint acts as a rheostat rather than a toggle switch [48].

This work highlights that accurate chromosome segregation results from the combined activities of error correction and the spindle assembly checkpoint [20] and suggests a simple procedure for determining how any particular condition impacts them to produce chromosome segregation errors:

1. *Measure the fraction of cells with the correct number of chromosomes after cell division:* This fraction, i.e. the fraction of cells with |∆*N* | = 0, is determined by the ratio of the checkpoint failure rate, *k*_*f*_, to the error correction rate, *k*_*b*_, with the relationship given by Eq. 3. If a perturbation causes the fraction of cells with the correct number of chromosomes to decrease, this could be caused by an increase in the checkpoint failure rate, a decrease in the error correction rate, or both.
2. *Measure the mean anaphase time:* A perturbation that decreases fraction of cells with the correct number of chromosomes, but increases mean anaphase times likely results from a decrease in the rate of error correction, i.e. Eq. 5 . A perturbation that decreases fraction of cells with the correct number of chromosomes, but decreases mean anaphase times likely results from an increase in the rate of checkpoint failure, i.e. Eq. 4 .

Note that these guidelines are heuristics that will not always be valid, and a more quantitative and rigorous understanding can be obtained by using the forced anaphase experiments, measurements of the distribution of spontaneous anaphase times and the distribution of spontaneous |∆*N* | values, and numerically fitting the results using the full state-dependent faulty checkpoint model to determine all model parameters. None the less, these guidelines are supported by our observations and calculations, and will lead to correct conclusions in many cases of interest. For example, RPE1 MAD2 RNAi cells and U2-OS monastrol cells have a similarly low fraction of cells with the correct number of chromosomes after cell division (58.8% and 53.7% respectively). However, the mean anaphase time of RPE1 MAD2 RNAi cells are very short (9.7 ± 0.3 min), while the mean anaphase time of U2-OS monastrol cells are very long (49.8 ± 2.1 min), consistent with RPE1 MAD2 RNAi cells having faulty checkpoint with properly functioning error correction and U2-OS monastrol cells having slightly misfunctioning checkpoint and greatly perturbed error correction (which is born out by use of forced anaphase experiments, measurements of the distribution of spontaneous anaphase times, and fitting with the full state-dependent faulty checkpoint model). This approach could be a useful tool when screening cell lines or perturbations – enabling estimates of error correction rates and checkpoint failure rates with relatively simple live cell imaging.

This work illustrates not only how error correction rates and checkpoint failure rates impact chromosome segregation errors, but also how they impact mitotic duration: more rapid cell division can be achieved by increasing either rate. Therefore, evolutionary pressure to increase the speed of mitosis might select for increased checkpoint failure rates as a byproduct. This may provide an explanation for the weak spindle assembly checkpoint and high rate of mitotic errors in many rapidly dividing early embryos, including human, mouse, zebrafish and Xenopus [49–53]. Such an outcome is reminiscent of the trade-off between risk and speed which has been proposed to be a fundamental feature of all biological checkpoints [54].

## Materials and Methods

### Cell lines

A stable hTERT RPE-1 cell line expressing CENP-A::sfGFP, mCherry::alpha tubulin, and emiRFP 670::hCentrin2 was used for all RPE-1 experiments. A stable U2-OS cell line expressing CENP-A::sfGFP, mCherry::alpha tubulin, and emiRFP 670::hCentrin2 was generated using a lentiviral system (Effectene Transfection Reagent, Qiagen) and selected using puromycin, blasticidin, and neomycin. Cells were further selected using fluorescence-activated cell sorting to eliminate CENP-A overexpression.

Both cell lines were maintained in Dulbecco ‘s modified Eagle ‘s medium GlutaMAX (DMEM GlutaMAX, Thermo Fisher) supplemented with 10% Fetal Bovine Serum (FBS, Thermo Fisher) and 50 IU ml^−1^ penicillin and 50 ug ml^−1^ streptomycin (Thermo Fisher) at 37°C in a humidified atmosphere with 5% CO2. Cells were regularly validated as mycoplasma free by a PCR-based mycoplasma detection kit (Southern Biotech).

### Live-cell imaging

All live-cell spinning-disk confocal microscopy imaging was performed as follows. Cells were plated on 25 mm diameter, #1-thickness, round coverglasses (Bioscience Tools) in 35mm dishes to 70-80% confluency (control and UMK57) or 30-40% confluency (monastrol washout) the day before experiments.

For unsynchronized hTERT RPE-1 samples, media was replaced with DMEM containing 1:4000 SPY650-DNA (Cytoskeleton) at least one hour before imaging. For unsynchronized U2-OS samples, media was replaced with DMEM containing 1:2000 SPY650-DNA (Cytoskeleton) at least one hour before imaging.

For the monastrol washout samples, mitotic cells were shaken off of the coverslips and the DMEM was replaced with 100µM monastrol (SelleckChem) DMEM. The samples were imaged after 2 hours (RPE-1) or 5 hours (U2-OS) of incubation in drug media.

Imaging experiments were performed on a home-built spinning disk confocal microscope (Nikon Ti2000, Yokugawa CSU-X1) with 488nm, 561nm, and 647nm lasers, a CMOS camera (Hamamatsu) and a 60x oil immersion objective. Imaging was controlled using a custom MicroManager program. The samples were transferred to a custom-built cell-heater calibrated to 37°C (Bioscience Tools).

For unsynchronized samples, cells were covered with 750µL of Fluorobrite DMEM (Thermo Fisher) supplemented with 10mM HEPES (Thermo Fisher) and 1µM UMK57 (if relevant; ChemFarm) and covered with 2mL of mineral oil. Up to 22 cells with condensed chromosomes (selected based on SPY650-DNA signal) were imaged in timelapses. Three separate fluorescence channels were acquired every 1 minute with 50ms exposure for 488nm, 546nm, and 647nm excitation in 3 z planes with 3µm spacing. End of nuclear envelope breakdown time was determined based on movies – for cells that underwent NEBD before the start of imaging, NEBD end times were backtracked using spindle morphologies.

For monastrol washout samples, cells were covered with 750µL of Fluorobrite DMEM supplemented with 10 mM HEPES and 100µM monastrol. Up to 45 mitotic cell positions were imaged in timelapses. Two separate fluorescence channels were acquired every 1 minute with 50ms exposure for both 488nm and 546nm excitation in 3 z planes. Monastrol was washed out after 2 minutes of imaging (2×1mL washes) and replaced with 750µl imaging media containing either drug or 0.5% v/v DMSO and covered with mineral oil. Drug concentrations used were: 10 µM TAK-981 (MedChemExpress), 1 µM UMK57 (ChemFarm), 30nM barasertib, 30nM volasertib, and 1 µM 5-ITu (SelleckChem).

For forcing anaphase, 750µL of 50µM AZ-3146 (SelleckChem) imaging media (with or without drug or DMSO) was added to the samples at the indicated times, for a final concentration of 25 µM AZ-3146. Anaphase times were recorded based on the time when kinetochores began separating in the timelapse movies. After all cells divided, high-resolution images were taken of the divided kinetochores for control and UMK57 samples, and both kinetochores and poles for monastrol washout samples (50ms exposure at full laser power for both 488nm and 647nm, 0.5µm z spacing) for at least 3 timepoints, spaced 10 minutes apart.

### Quantitative analysis of kinetochore count data

Z-stacks were run through the custom Python 3 kinetochore counting code in JupyterLab and manually adjusted for final kinetochore counts as previously described [17]. Kinetochore counting code can be found on Github (https://github.com/gloriaha/kinetocounter).

### Fitting of Mad2 RNAi data

For the simultaneous fitting of the spontaneous |∆*N* |*/*2 and spontaneous anaphase times, we prepared the data and fitting procedures as follows:

For the spontaneous |∆*N* |*/*2 and spontaneous anaphase times, we compared the data to the numerically simulated probability distributions of |∆*N* |*/*2 and anaphase onset times, as described in Appendix B and C. We will refer to the numerically predicted probability distribution of anaphase onset times as *P*_ana_ and the numerically predicted probability distribution of |∆*N* |*/*2 as *P*_|∆*N*|_.

For the spontaneous anaphase time data, timelapse movies with 1 minute resolution were used to estimate the anaphase onset time relative to NEBD end. The first frame where the sister kinetochores started moving apart was taken to be the anaphase onset time. Anaphase time data was divided into 1 minute bins and each bin was normalized by the total number of cells. The resulting residual expression was *𝒴*_hist,*i*_ − *P*_ana_(*𝒳*_hist,*i*_), where *𝒴*_hist,*i*_ was the fraction of cells in the bin starting at time *𝒳*_hist,*i*_.

We divided the spontaneous |∆*N* |*/*2 data into bins of size 1 (|∆*N* |*/*2 = 0, 1, 2…) from |∆*N* |*/*2 = 0 to |∆*N* |*/*2 = 46 and normalized each bin by the total number of cells. The resulting residual expression was *𝒴*_hist,*i*_ − *P*_|∆*N*|_(*𝒳*_hist,*i*_), where *𝒴*_hist,*i*_ was the fraction of cells in the bin starting at |∆*N* |*/*2 = *𝒳*_hist,*i*_.

We used the *k*_*b*_, *k*_*e*_, *C*_*E*,init_ from fitting the anaphase onset time and forced anaphase kinetochore count data from RPE-1 control cells without Mad2 RNAi (using the model allowing for statistical asymmetry and finite error rate, fit results taken from previous paper [17]: *k*_*b*_ = 0.57 ± 0.03 min^−1^, *k*_*e*_ = 0.001 ± 0.0003 min^−1^, *C*_*E*,init_ = 26 ± 9). We fixed these parameter values and simultaneously fit the anaphase onset time and |∆*N* |*/*2 distributions to find the best fit *t*_offset_ and *k*_*f*_ . For the state-dependent model, we fixed *p*_*f*_ = 0.5 after comparing the fit results with different values of *p*_*f*_ .

The simultaneous fit was done using a joint squared residual expression, with the numerically simulated probability distributions of the slowest first passage time and kinetochore count differences (distributions described in the Appendix):

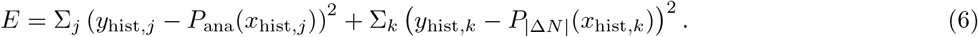

We chose to equally weight the anaphase time and kinetochore count data – some experimental conditions had more of one type of data than the other, so in order to equally consider the datasets in the fit, we normalized each by the total number of data points. We observed that skewing the weighting towards anaphase times or kinetochore counts led to poor fits of the other dataset. The best fit *t*_offset_ and *k*_*f*_ were determined using Scipy ‘s curve fit function on the summed squared residuals. We used the resulting covariance matrix to estimate the parameter fit errors using Numpy ‘s uncertainty module.

### Simultaneous fitting of data with full state-dependent SAC model (allowing for statistical asymmetric initial attachments, finite *k*_*e*_, and state-dependent model for faulty SAC)

For the simultaneous fitting of the spontaneous |∆*N* |*/*2, spontaneous anaphase times, and forced anaphase datasets, we prepared the data and fitting procedures as follows:

For fitting the forced anaphase data of the unsynchronized samples (not monastrol washout), cells were binned into two minute intervals, and means and standard deviations of the squared kinetochore count differences were calculated for each bin. For fitting the monastrol washout samples, we calculated the means and standard deviations of the squared kinetochore count differences for every timepoint (5 minute intervals from 5 to 30 minutes).

We used the following equation for the forced anaphase data predictions, which allowed for statistical asymmetry (derivation in Appendix of [17]):

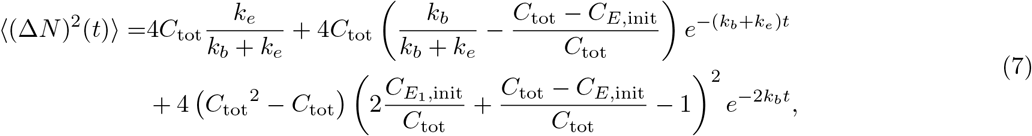

where 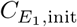 represents the average number of erroneous attachments at one pole. We constrained the parameters such that 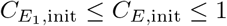.

The spontaneous |∆*N* |*/*2 and spontaneous anaphase times was prepared as described in the previous section.

The simultaneous fit was done using a joint squared residual expression, with the expression valid for statistically asymmetric initially erroneous attachments and the numerically simulated probability distribution of the slowest first passage time with a finite *k*_*e*_ and state-dependent model for a faulty checkpoint.

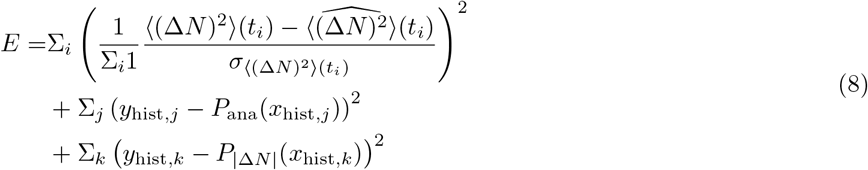

The best fit *k*_*b*_, *k*_*e*_, *C*_*E*,init_, *t*_offset_, 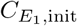, and *p*_*f*_ were determined using Scipy ‘s curve fit function on the summed squared residuals. We used the resulting covariance matrix to estimate the parameter fit errors and 2*σ* confidence interval (shaded region of plot) using Numpy ‘s uncertainty module.

## Acknowledgments

We thank Timothy Mitchison, Iain Cheeseman, Vinothan Manoharan, Alexey Khodjakov, and Andrea Musacchio for helpful discussions, Paul Dieterle and Hao Shen for helping develop the originally published modeling and analysis pipeline [17], and the Bauer Flow Cytometry Core for cell sorting. G.H. was supported by the QuantBio Student Fellowship through the NSF-Simons Center for Mathematical and Statistical Analysis of Biology at Harvard (1764269). This work was funded by NSF award DBI-1919834. Research reported in this publication was supported by an award from the Zuckerman Travel and Research STEM Fund at Harvard University. A.A. thanks support from the Clore Center for Biological Physics. We also acknowledge support from the CCBX program of the Center for Computational Biology of the Flatiron Institute.

