## Appendix for "Contributions of error correction and the spindle assembly checkpoint to mitotic timing and fidelity"

### A. Numerics for the erroneous chromosome segregation model

Since we have $C_{\rm tot}$ pairs of chromosomes, we treat the system as a Markov chain with $(C_{\rm tot}+1)$ states of 0, 1, 2, ..., $C_{\rm tot}$ incorrect pairs. For each pair of chromosomes, it can go from the incorrect state to the correct state at rate $k_b$, and from the correct state to the incorrect state at rate $k_e$. We define the transition probabilities $P_i(t) = \Pr\left(X(t) = i | X(0) = j\right)$, which denotes the probability for the system to start with a given initial condition state $j$ and transit to state $i$ by time $t$. We consider a sloppy check-point system so that at each state $i$ there is a rate $f_i$ to divide and exit the system. For the constant rate model, $f_i = k_f$, while for the state-dependent model, $f_i = k_f p_f^{i-1}$. The master equation is then:

$$\frac{dP_1}{dt} = -[(C_{\rm tot}-1)k_e + k_b]P_1 + 2k_bP_2 - f_1P_1 \tag{1}$$

$$\frac{dP_i}{dt} = (C_{\rm tot}-i+1)k_eP_{i-1} - [(C_{\rm tot}-i)k_e + ik_b]P_i + (i+1)k_bP_{i+1} - f_iP_i \quad (i = 2, ..., C_{\rm tot}-1) \tag{2}$$

$$\frac{dP_{C_{\rm tot}}}{dt} = k_eP_{C_{\rm tot}-1} - Nk_bP_{C_{\rm tot}} - f_{C_{\rm tot}}P_{C_{\rm tot}} \ . \tag{3}$$

We did not specify the equation for $P_0$ here since all the pairs being correct would lead to division directly (i.e. it is the absorbing state). We can numerically solve this set of equations and calculate the probability density of dividing at certain state $i$ at time $t$: $f_iP_i(t)$. We can integrate the probability density over a long enough period of time to calculate the probability for the system to divide at certain state $i$:

$$P_{div,i} = \int_{t=0}^{\infty} f_iP_i(t)dt \ . \tag{4}$$

To get the $|\Delta N|$ distribution, we consider a binomial model in which each incorrectly divided pair goes to left or right cell randomly. Then the $|\Delta N|/2$ distribution is given by:

$$P(|\Delta N|/2 = |i-2j|) = \sum_{i=0, 0\le j\le i}^{C_{\rm tot}} \binom{i}{j} 2^{-i} P_{div,i} \ . \tag{5}$$

To get the probability distribution of dividing at time $t$, we simply notice that the probability of the divided and undivided states should add up to one. Thus we have the anaphase onset time distribution:

$$P_{\rm anaphase\,onset\,time}(t) = \frac{d}{dt}(1 - \sum_{i=1}^{C_{\rm tot}} P_i) = -\sum_{i=1}^{C_{\rm tot}} \frac{dP_i}{dt} \ . \tag{6}$$

Equivalently we can express it by adding up the probability density for dividing at each state $i$:

$$P_{\rm anaphase\,onset\,time}(t) = k_bP_1 + \sum_{i=1}^{C_{\rm tot}} f_iP_i \ . \tag{7}$$

### B. Last step approximation of the mean anaphase time

We can derive an approximate expression for the mean anaphase time within this model. In calibrating the change in mean anaphase time due to faulty spindle assembly checkpoint, we hypothesize that the major contribution comes from the last step going from the state 1 to division.

We begin by noting that the mean first passage time to reach division is the sum of mean first passage time to reach state 1 and the expected time to go from state 1 to division. When there is no spindle assembly checkpoint failure, i.e., $k_f = 0$, the expected time to go from state 1 to division is $1/k_b$. When there is a nonzero rate $k_f$ for the system to divide at state 1, the total rate for the system to go from state 1 to division is $k_b + k_f$, and the expected time becomes $1/(k_b + k_f)$. Thus, the difference in the mean anaphase time induced by faulty spindle assembly checkpoint in the last step is simply $1/(k_b + k_f) - 1/k_b$. Under this approximation of considering only the last step going from state 1 to division, we have:

$$\langle t_{\text{ana}} \rangle \approx \langle t_{\text{ana}} \rangle_{k_f=0} + \left( \frac{1}{k_b + k_f} - \frac{1}{k_b} \right) \approx \frac{\gamma + \log(C_{E.\text{init}})}{k_b} + t_{\text{offset}} + \left( \frac{1}{k_b + k_f} - \frac{1}{k_b} \right) . \tag{8}$$

where $\gamma$ is Euler's constant. For the second step in the equation we express the mean anaphase time for $k_f = 0$ under the approximation of very small error rate $k_e \approx 0$.

### C. Last step approximation of the fraction of cells with $|\Delta N|/2 = 0$

Similarly, we can express the fraction of cells that has $|\Delta N|/2 = 0$ within this last step approximation:

$$P(|\Delta N| = 0) \approx \frac{k_b}{k_b + k_f} . \tag{9}$$

This would be an overestimation of the fraction of cells that have $|\Delta N|/2 = 0$ since we neglect the possibility of dividing at states other than state 1.

TABLE I: Simultaneous state-dependent SAC model fit results for RPE-1 monastrol washout. Concentrations used: 0.5% v/v DMSO, 30 nM volasertib (Plk1i), 10 µM TAK-981 (SUMOi), 1 µM UMK57, 50 nM alisertib (AurAi), 1 µM 5-ITu (Haspin i), 30 nM barasertib (AurBi)

| Condition | $k_b$ (min$^{-1}$) | $k_e$ (min$^{-1}$) | $C_{E,\text{init}}$ | $t_{\text{offset}}$(min) | $A_0$ | $k_f$(min$^{-1}$) |
|---|---|---|---|---|---|---|
| **DMSO control** | $0.15 \pm 0.02$ | $0 \pm 0.001$ | $46 \pm 26$ | $6 \pm 2$ | $620 \pm 100$ | $0.005 \pm 0.001$ |
| **Plk1 inhibitor** | $0.15 \pm 0.01$ | $0 \pm 0.001$ | $37 \pm 19$ | $3 \pm 2$ | $1240 \pm 220$ | $0.05 \pm 0.004$ |
| **SUMO inhibitor** | $0.15 \pm 0.01$ | $0 \pm 0.001$ | $46 \pm 24$ | $6 \pm 2$ | $680 \pm 130$ | $0.04 \pm 0.004$ |
| **UMK57 (MCAK activator)** | $0.09 \pm 0.02$ | $0 \pm 0.001$ | $46 \pm 60$ | $0 \pm 8$ | $230 \pm 50$ | $0.002 \pm 0.001$ |
| **Aurora A inhibitor** | $0.14 \pm 0.01$ | $0 \pm 0.001$ | $46 \pm 9$ | $5 \pm 2$ | $0 \pm 23$ | $0.04 \pm 0.004$ |
| **Haspin inhibitor** | $0.11 \pm 0.02$ | $0 \pm 0.001$ | $30 \pm 70$ | $0 \pm 17$ | $1110 \pm 170$ | $0.05 \pm 0.008$ |
| **Aurora B inhibitor** | $0.16 \pm 0.03$ | $0.002 \pm 0.002$ | $46 \pm 33$ | $6 \pm 2$ | $90 \pm 60$ | $0.04 \pm 0.008$ |

TABLE II: Simultaneous state-dependent SAC model fit results for RPE-1 and U2-OS cells

| Condition | $k_b$ (min$^{-1}$) | $k_e$ (min$^{-1}$) | $C_{E,\text{init}}$ | $t_{\text{offset}}$(min) | $A_0$ | $k_f$ (min$^{-1}$) |
|---|---|---|---|---|---|---|
| **RPE-1 control** | $0.57 \pm 0.02$ | $0.001 \pm 0.0002$ | $26 \pm 5$ | $10.8 \pm 0.3$ | $0 \pm 60$ | $0.002 \pm 0.003$ |
| **RPE-1** 1 µM **UMK57** | $0.38 \pm 0.04$ | $0.007 \pm 0.003$ | $42 \pm 12$ | $7.8 \pm 0.5$ | $0 \pm 130$ | $0.008 \pm 0.002$ |
| **U2-OS control** | $0.18 \pm 0.01$ | $0 \pm 0.0003$ | $11 \pm 4$ | $10 \pm 2$ | $128 \pm 20$ | $0.06 \pm 0.004$ |
| **U2-OS** 100 nM **UMK57** | $0.17 \pm 0.001$ | $0 \pm 0.0001$ | $8 \pm 0.03$ | $13 \pm 0.1$ | $68 \pm 9$ | $0.08 \pm 0.004$ |
| **U2-OS monastrol washout DMSO** | $0.06 \pm 0.01$ | $0 \pm 0.0002$ | $20 \pm 50$ | $0 \pm 34$ | $550 \pm 80$ | $0.04 \pm 0.004$ |

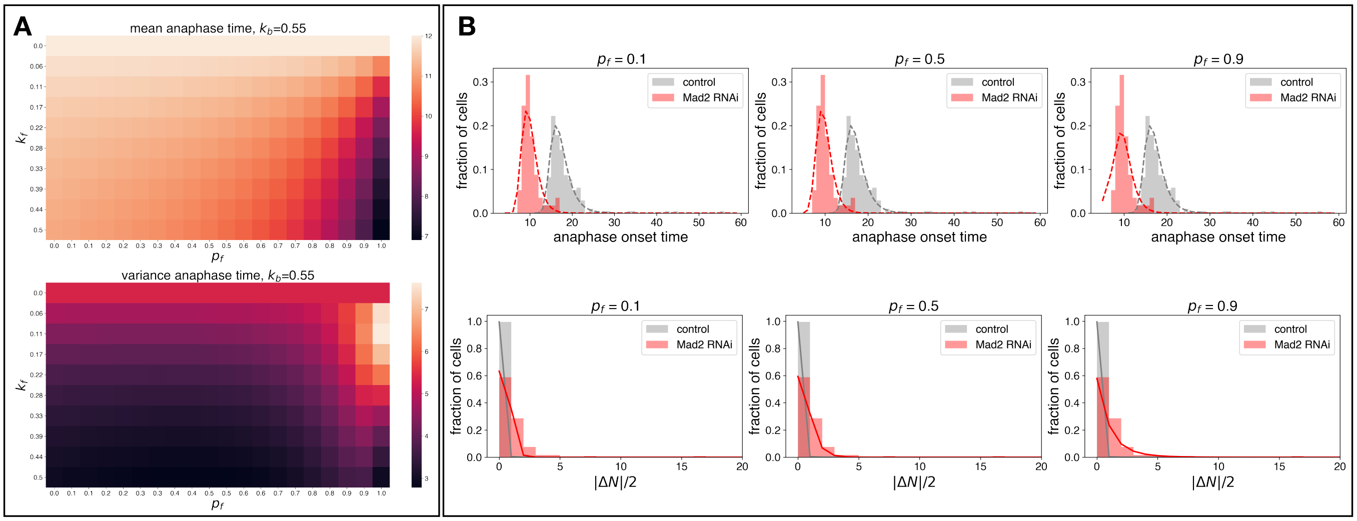


FIG. 1: **Dependence of anaphase time and kinetochore count difference distribution on** $p_f$. (A) State-dependent SAC model depdence of mean and variance of anaphase time distribution on $p_f$ and $k_f$. Both mean and variance have a stronger dependence on $k_f$ than $p_f$ for a reasonable range of $p_f$. (B) Fit results of control vs. Mad2 RNAi data while fixing $p_f$. When $p_f$ is too low ($p_f = 0.1$), the kinetochore count distribution is under-predicted, while when $p_f$ is too high ($p_f = 0.9$), the anaphase time distribution is poorly predicted and the kinetochore count difference distribution is slightly over-predicted.

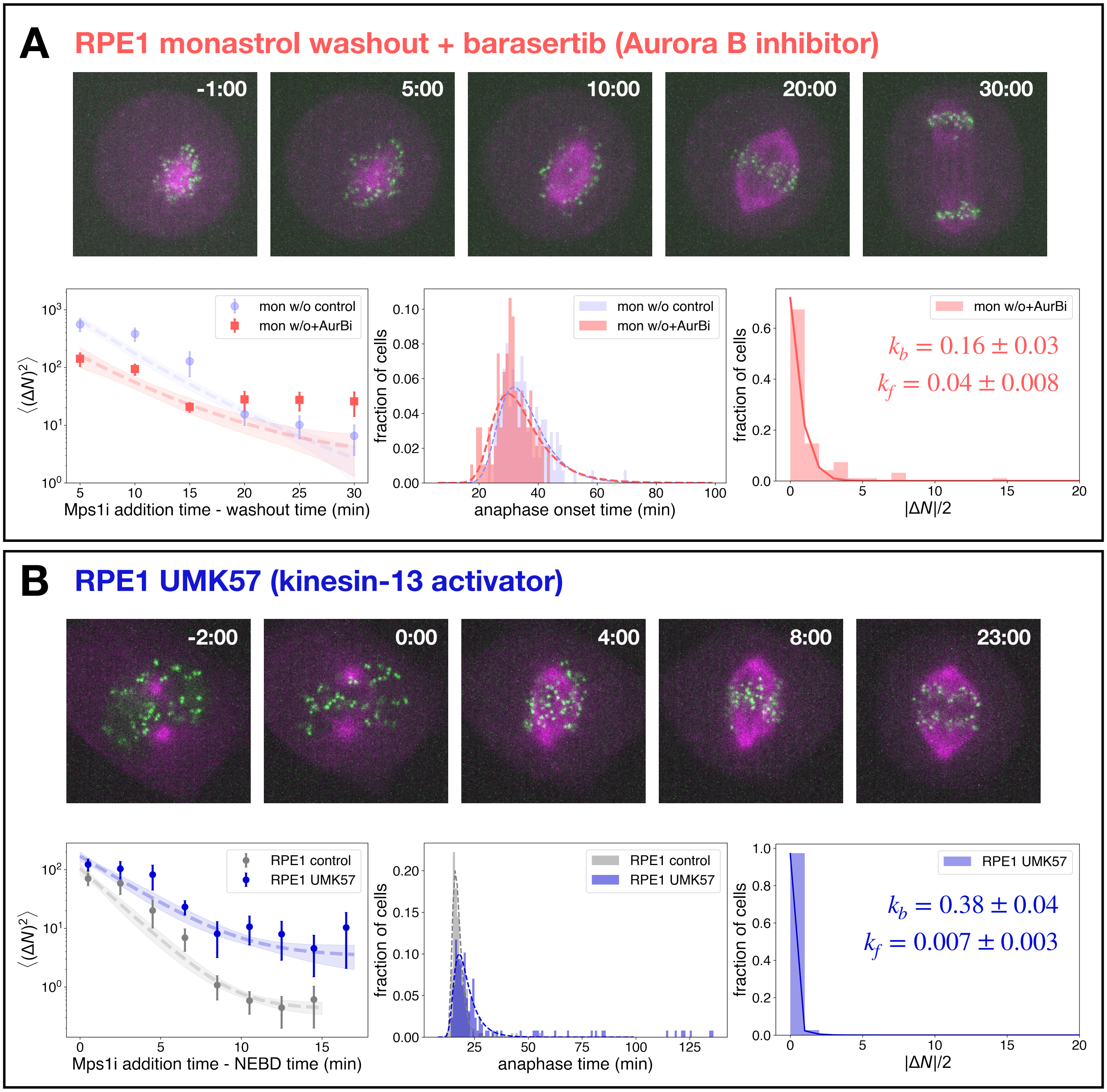


FIG. 2: **Results for RPE1 monastrol washout with Aurora B inhibition (A) and for RPE1 with kinesin-13 activation (B)**. (A) Example movie snapshots during RPE1 monastrol washout with 30nM barasertib, forced anaphase assay, spontaneous anaphase time distribution, and spontaneous kinetochore count difference distributions (B) Example movie snapshots during spindle assembly of RPE1 with 1 µM UMK57 (kinesin-13 potentiator), forced anaphase assay, spontaneous anaphase time distribution, and spontaneous kinetochore count difference distribution.
